# Cytoplasmic mRNA decay factors modulate RNA polymerase II processivity in 5’ and 3’ gene regions in yeast

**DOI:** 10.1101/741058

**Authors:** Jonathan Fischer, Yun S. Song, Nir Yosef, Julia di Iulio, L. Stirling Churchman, Mordechai Choder

## Abstract

mRNA levels are determined by the balance between mRNA synthesis and decay. Factors that mediate both processes, including the 5’ to 3’ exonuclease Xrn1, are responsible for the cross talk between the two processes in a manner that buffers steady-state mRNA levels. However, these proteins’ roles in transcription remain elusive and controversial. Applying NET-seq to yeast cells, we show that Xrn1 functions mainly as a transcriptional activator and that its disruption manifests via the reduction of RNA polymerase II (Pol II) occupancy downstream of transcription start sites. We combine our data and novel mathematical modeling of transcription to suggest that transcription initiation and elongation of targeted genes is modulated by Xrn1. Furthermore, Pol II occupancy markedly increases near cleavage and polyadenylation sites in *xrn*1Δ cells while its activity decreases, a characteristic feature of backtracked Pol II. We also provide indirect evidence that Xrn1 is involved in transcription termination downstream of polyadenylation sites. Two additional decay factors, Dhh1 and Lsm1, seem to function similarly to Xrn1 in transcription, perhaps as a complex, while the decay factors Ccr4 and Rpb4 also perturb transcription in other ways. Interestingly, DFs are capable of differentiating between SAGA- and TFIID-dominated promoters. These two classes of genes respond differently to *XRN*1 deletion in mRNA synthesis and differentially utilize mRNA decay pathways, raising the possibility that one distinction between the two types of genes lies in the mechanism(s) that balance these processes.

## Introduction

Steady-state mRNA levels are determined by the balance between synthesis and decay rates. Once thought to function separately, recent studies have discovered that these two processes are linked. In previous work we showed that the major cytoplasmic yeast mRNA degradation pathway, consisting of the decapping enzyme Dcp1/2, the decapping activator Pat1/Lsm1-7, the helicase Dhh1, and the 5’-3’ exonuclease Xrn1, shuttles between the cytoplasm and the nucleus to participate in both processes. Notably, the elements of this pathway were found to degrade most mRNAs in the cytoplasm while stimulating transcription in the nucleus. The proteins Dcp2, Lsm1, and Xrn1 were further shown to bind chromatin, probably as a complex, and to stimulate transcription initiation and elongation (Haimovich et al. 2013). We also uncovered a connection between how Xrn1 functions in transcription and mRNA decay by revealing the correlation between the effects of Xrn1 disruption on mRNA synthesis and decay in the nucleus and cytoplasm, respectively (Haimovich et al. 2013; Medina et al. 2014). We subsequently ranked genes according to their responsiveness to Xrn1 disruption in optimally proliferating yeast cells; the most responsive were dubbed the “Xrn1 synthegradon” and consisted of genes whose transcription and decay rates exhibited the highest sensitivity to Xrn1 disruption (Medina et al. 2014). This group is highly enriched with genes required for cell growth and proliferation, including genes encoding ribosome biogenesis and translation factors.

“Classical” mRNA decay factors are not the only bridges between transcription and mRNA decay. For example, Rpb4 and Rpb7, two canonical RNA polymerase II (Pol II) subunits, function in both processes (Choder 2004; Goler-Baron et al. 2008; Lotan et al. 2005; Lotan et al. 2007; Schulz et al. 2014; Shalem et al. 2011), and even promoters are capable of regulating mRNA decay (Bregman et al. 2011; Trcek et al. 2011). Hence the cross talk between mRNA synthesis and decay is complex and involves an interplay between canonical transcription and degradation factors. Although the links are clear, the mechanism mediating mRNA buffering remains enigmatic and controversial. Some publications have proposed a simple feedback mechanism involving a repressor (Sun et al. 2012; Sun et al. 2013), though others have suggested that components of the mRNA decay machinery function directly in transcription (Haimovich et al. 2013; Medina et al. 2014). In fact, the former articles proposed that the deletion of Xrn1 leads to transcription activation whereas the latter group asserted the opposite.

The realization of the critical role of mRNA buffering requires changes in the approaches used to analyze transcription. In the past, mRNA levels were regarded as a good proxy for transcription, and prior studies have relied upon changes in these levels to infer alterations in transcription. As an example, earlier work classified genes as SAGA- or TFIID-dominated based on measured changes in mRNA levels after inactivation of central components of the SAGA (mainly Spt3) or TFIID (mainly Taf1) complexes (Basehoar et al. 2004; Huisinga and Pugh 2004). However, it was recently reported that virtually all promoters recruit both the SAGA and TFIID complexes and recent transcriptional profiling experiments demonstrated that mutations in either complex result in widespread defective transcription (Baptista et al. 2017; Bonnet et al. 2014; Warfield et al. 2017). Nonetheless, the disruption of most components of either complex did not lead to decreases in the levels of most mRNAs due to feedback mechanisms that involve mRNA decay (Baptista et al. 2017; Bonnet et al. 2014; Warfield et al. 2017). It is now clear that mRNA levels are not simply determined by two unrelated processes of mRNA synthesis and decay; rather, each of these processes affects the other by a hitherto elusive mechanism.

Following transcription initiation, many metazoan genes undergo a regulatory step termed promoter-proximal pausing (reviewed recently in Chen et al. 2018 and Wissink et al. 2019). Specifically, after transcribing 30–120 nucleotides downstream of transcription start sites (TSS), Pol II pauses; its release into productive elongation requires the activity of specific factors, including TFIIS. Following its release, Pol II interacts with additional elongation factors that modulate its processivity. It was recently reported that the release of the mammalian Pol II from a paused state in the promoter-proximal region is a key step in the regulation of transcription, both generally (Sheridan et al. 2019) and in response to environmental stress (Bartman et al. 2019; Sheridan et al. 2019). Interestingly, Pol II recruitment rate was proposed to have only a marginal impact on overall transcription rates (Bartman et al. 2019). Conversely, common wisdom posits that promoter-proximal pausing does not play a major role in budding yeast as it is less prominent than in metazoans (Adelman and Lis 2012). However, there is evidence that Pol II in *S. cerevisiae* accumulate downstream of TSS (Churchman and Weissman 2011), though this phenomenon and its contribution to transcriptional regulation has been little-studied. In contrast with the inconclusive nature of 5’ pausing, a conspicuous Pol II pausing event does occur at polyadenylation sites (PAS) (e.g., Mayer et al. 2017). It is plausible that this pausing is required to provide the necessary time for the assembly of polyadenylation (PA) machinery, but gaps in the mechanistic understanding of this pausing event persist. Nevertheless, it is clear that factors of the PA pathway affect transcription termination events that occur downstream.

To probe the effects of mRNA decay factors (DFs) on transcription, we employed Native Elongating Transcript sequencing (NET-seq), an experimental protocol which assays Pol II occupancy at single nucleotide resolution. This technique sequences nascent RNA strands attached to actively engaged Pol II (Churchman and Weissman 2012) and maps the 3’ ends of nascent RNAs to yield the positions of Pol II active sites. Therefore, unlike RNA-seq, NET-seq data are not confounded by mRNA decay rates and give the precise locations of bound Pol II. Additionally, non-coding RNAs (ncRNAs) are frequently difficult to detect using RNA-seq because of their low transcript stabilities, but are easily identified using NET-seq, permitting more thorough investigations of additional classes of transcripts. In contrast to other transcription profiling methods such as Genomic Run-On (GRO) (García-Martínez et al. 2004) and its high-resolution cousin Biotin Genomic Run-On (BioGRO) (Jordán-Pla et al. 2014; Jordán-Pla et al. 2016), which assay only actively elongating Pol II, NET-seq can report both elongating Pol II as well as arrested Pol II (Churchman and Weissman 2011). As a result, run-on methods and NET-seq are particularly informative when used in combination and can reveal information about Pol II processivity and pausing.

In light of the poorly understood mechanisms linking mRNA synthesis and decay, we applied NET-seq to obtain Pol II occupancy profiles in various DF deletion strains to facilitate study of the roles of DFs in transcription. In addition to effects on initiation, Xrn1 and our other studied DFs seem to affect transcription primarily via elongation changes which are plausibly attributable to modified Pol II pausing and backtracking. These effects primarily manifest in the ends of genes, occurring ∼100 bp downstream of TSS in the 5’ end and extending from PAS roughly 75 bp into the 3’ ends of gene bodies. Similar changes in Pol II occupancy were identified in the 5’, but not 3’, ends of ncRNAs, implicating DFs in the regulation of the early stages of ncRNA transcription. Furthermore, deletion of *XRN*1 affected Pol II elongation efficiency in a manner consistent with enhanced Pol II backtracking. We additionally employed a recently developed mathematical model (Erdmann-Pham et al. 2018) to infer changes in spatial transcriptional dynamics. This methodologically novel model uses our metagene profiles to estimate baseline values for relative initiation and elongation rates while offering a framework to systematically vary unknown parameters. This allowed us to perform *in silico* experiments suggesting that Xrn1 is required for efficient initiation of its target genes. In contrast to the most affected genes, NET-seq signals increase in response to Xrn1 disruption in a small repertoire of so-called “repressed” genes. Interestingly, these genes displayed demonstrably different 5’ and 3’ occupancy patterns upon DF deletion. Given these observed differences and comparisons with external data, we propose that the considered DFs modulate transcription of a subset of genes, perhaps as a complex, mainly via the regulation of pausing and backtracking during the early and late stages of transcription.

## Results

### Deletions of mRNA decay factors lead to overall decreases in Pol II occupancy

Previously we reported that Xrn1 binds to promoters and gene bodies and directly stimulates transcription initiation and elongation (Haimovich et al. 2013). Shortly thereafter, Sun et al. reported that the deletion of *XRN*1 leads to the upregulation of transcription, implying that Xrn1 represses transcription (Sun et al. 2013). To resolve this discrepancy and gain insight into the mechanisms linking transcription and mRNA decay, we used NET-seq to compare Pol II occupancy in wild type (WT) strains and those carrying a deletion of *XRN*1 (*xrn*1Δ). Overall Pol II occupancy in *xrn*1Δ cells generally decreased (Fig. 1A), indicating the downregulation of transcription. Moreover, the fold change distribution had a heavier negative than positive tail, highlighting that a subset of genes was affected more than others. Our results indicate that Pol II occupancy is negatively affected by *XRN*1 deletion, consistent with its proposed role as a transcriptional activator (Haimovich et al. 2013; Medina et al. 2014). We further studied additional mutant strains, each carrying a deletion of either *CCR*4, *DHH*1, or *LSM*1, and observed decreases in Pol II occupancy in each respective knockout, although not as strongly as those found in *xrn*1Δ cells (Fig. S1A). These results support roles for the encoded proteins as transcriptional stimulators.

**Fig. 1.**
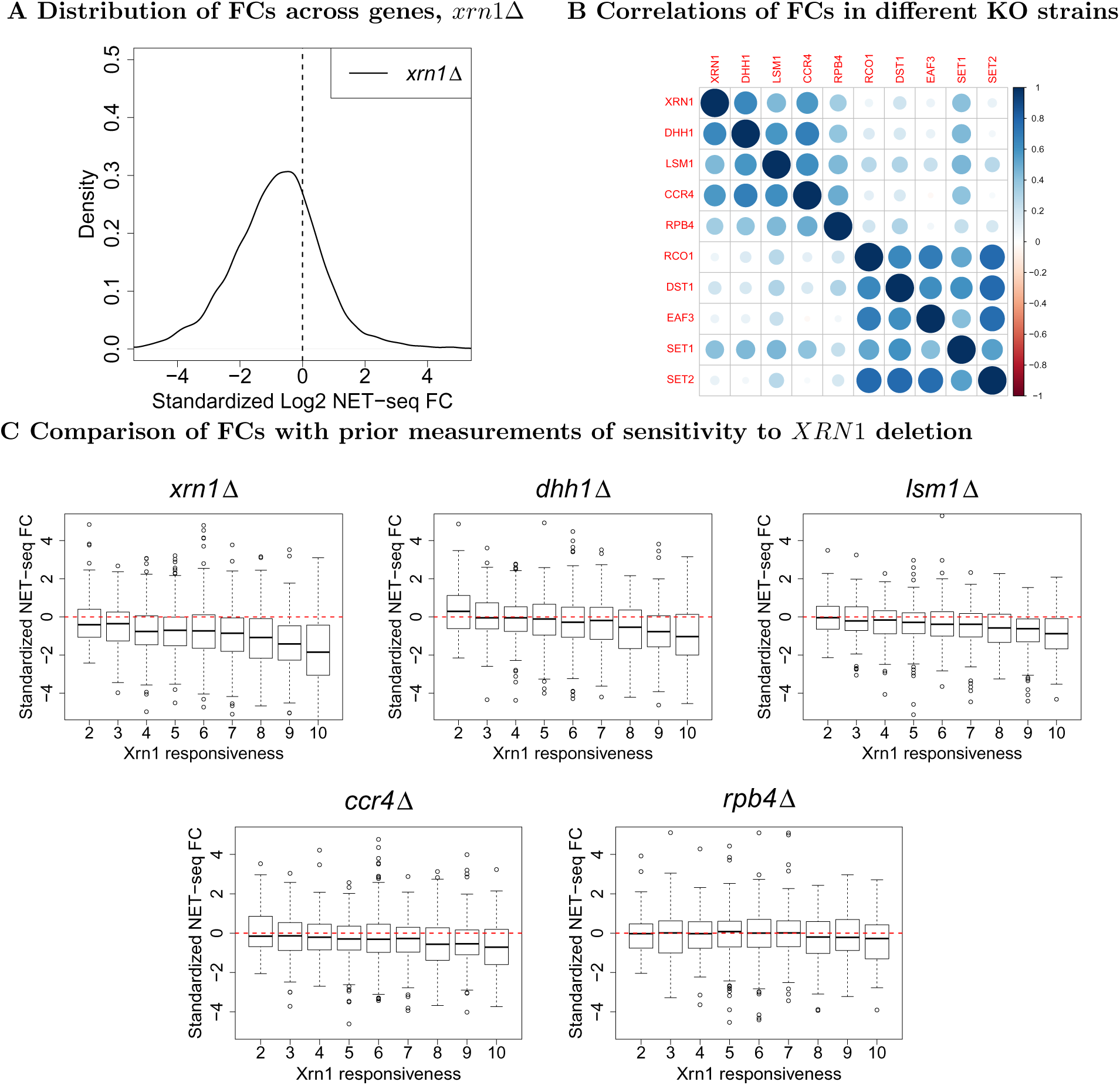
Fold changes in NET-seq Pol II occupancy. (A) NET-seq reads were aggregated within annotated gene boundaries (TSS to PAS) and applied DESeq2 (Love et al. 2014) to estimate standardized fold changes in each gene’s normalized signal with respect to the WT. Both *xrn*1Δ and WT were done in two replicates. (B) The visualized correlation matrix for standardized fold changes in NET-seq reads in genes. Each entry corresponds to the Spearman correlation between the fold change with respect to the wild type in NET-seq reads in annotated genes. *xrn*1Δ, *dhh*1Δ, *lsm*1Δ, *ccr*4Δ, and *rpb*4Δ come from the experiments associated with this paper whereas the rest come from (Churchman and Weissman 2011). Fold changes were estimated using DESeq2 (Love et al. 2014) with both experiments analyzed simultaneously. **C**: Genes were stratified using previously obtained measures of Xrn1 responsiveness, an aggregated measure of the sensitivity of synthesis and decay rates to Xrn1 deletion as measured in (Medina et al. 2014). A value of 2 indicates the lowest sensitivity and 10 the highest. Standardized NET-seq fold changes from our experiments were then plotted for genes falling into each responsiveness classification.

In earlier work, we found that Xrn1, Lsm1, and Dcp2 produced highly similar ChIP-exo profiles (Haimovich et al. 2013), raising the possibility that they function in transcription as a complex. To examine potential complex associations among DFs, we sought to characterize the extent to which deletion strains induced similar genome-wide transcriptional responses. We constructed a correlation matrix of gene Pol II fold changes (Fig. 1B), uncovering strong similarities in transcriptomic response to deleted DFs. As an external control, we further examined the correlations between our NET-seq data and previously generated NET-seq profiles from strains carrying respective deletions of *EAF* 3, *RCO*1, and *SET*2, genes encoding a set of proteins which are known to act in concert and are required for proper function of the Rpd3S H4 deacetylation complex (Churchman and Weissman 2011). Strains carrying deletions in each of *DST*1 and *SET*1, encoding proteins that regulate Pol II release from backtracking and promoter direction-ality (Churchman and Weissman 2011), were also included. Fold changes among these non-DF strains correlated well (*ρ* ∼ 0.75 − 0.8) as expected due to their shared functions. In contrast, correlations between these non-DFs and our studied DFs were typically much lower (Fig. 1B), though this may be in part because they were collected in a different experiment. Nonetheless, the strong correlations among DF mutants suggest that they may act together in a complex as has been proposed for Lsm1 and Xrn1 (Haimovich et al. 2013). Rpb4 is a protein that functions in both mRNA synthesis and decay (Choder 2004; Duek et al. 2018; Lotan et al. 2005). To examine whether Rpb4 function is indeed related to those of our studied DFs, we performed NET-seq on an *rpb*4Δ strain and compared it to our other samples. As suspected, the *rpb*4Δ NET-seq profile correlated well with all considered DFs (*ρ* ∼ 0.5), attaining its highest correlation with *dhh*1Δ (Fig. 1B). Interestingly, *rpb*4Δ NET-seq data correlated better with the studied DF KOs than with KOs of factors that function in transcription, e.g. *dst*1Δ (Fig. 1B). This pattern of correlations suggests that Rpb4 functions similarly to the studied DFs in linking mRNA synthesis and decay, consistent with its known interactions with both Pol II (Choder 2004) and the scaffold of the mRNA decay complex, Pat1 (Lotan et al. 2005), as well as its distinct function in transcription and in the major cytoplasmic mRNA decay pathway (Duek et al. 2018; Goler-Baron et al. 2008; Lotan et al. 2005; Lotan et al. 2007; Shalem et al. 2011). We therefore included the *rpb*4Δ strain in our subsequent analyses.

To compare results obtained via NET-seq to other RNA quantification methods, we reviewed publicly available data for knockouts which were considered in our experiment. We obtained RNA-seq fold changes for both *dhh*1Δ and *lsm*1Δ and GRO data for *rpb*4Δ and *xrn*1Δ (He et al. 2018; García-Martínez et al. 2015; Gutiérrez et al. 2017; Haimovich et al. 2013). Comparisons of NET-seq, RNA-seq, and GRO data convincingly demonstrated that reads correlated much more strongly by protocol than by condition for both raw reads (not shown) and fold changes (Fig. S2). This highlights the fact that each protocol reports different aspects of gene expression; for example, whereas NET-seq reports Pol II occupancy, GRO reports Pol II elongation activity, and RNA-seq captures steady-state RNA levels. The low correlations of fold changes among different quantification methods further demonstrates the importance of the cross talk between mRNA synthesis and decay.

To examine whether Xrn1 or Rpb4 are required for the overall processivity of Pol II, we compared NET-seq signals with previously reported GRO signals (García-Martínez et al. 2015; Haimovich et al. 2013). GRO results are sensitive to backtracking because the RNA 3’ end of backtracked Pol II is displaced from the active site and transcription elongation cannot proceed *in vitro*. The log2 ratio between GRO signal and Pol II occupancy detected by NET-seq (henceforth the elongation efficiency) is substantially compromised in the *xrn*1Δ strain (Fig. S1B), indicating that Xrn1 helps prevent or resolve backtracking, thus mediating proper elongation of Pol II. Correspondingly, we consistently found no correlation between NET-seq and GRO data (Fig. S2). In comparison to *xrn*1Δ, the *rpb*4Δ strain displayed a smaller decrease in efficiency, so it may not be as important as Xrn1 in the prevention or resolution of backtracking across the genome. However, as we show later, Rpb4 does impact Pol II activity in the 5’ and 3’ ends of genes. To further probe the impact of Xrn1 and Rpb4 on elongation, we compared fold changes in elongation efficiency to gene length. Consistent with the above, fold changes in *xrn*1Δ were more negative than those in *rpb*4Δ for genes of all lengths, but both deletion strains showed that longer genes tended to see larger reductions in elongation efficiency (Fig. S3). These findings suggest Xrn1 and, to a lesser extent, Rpb4 are important for efficient Pol II elongation in a manner which becomes more essential for longer genes (Medina et al. 2014; Verma-Gaur et al. 2008). Importantly, we do not suggest that Rpb4 is not important for transcription, as was demonstrated previously by other means (see Introduction); rather, we conclude that the overall effects of *RPB*4 deletion identified using GRO are similarly reflected in the NET-seq data.

We previously rated genes according to the sensitivity of their mRNA synthesis and decay rates to *XRN*1 deletion. mRNAs whose synthesis and decay were highly responsive to Xrn1 disruption were named the “Xrn1 synthegradon”, while those least affected were dubbed the “Xrn1 anti-synthegradon” (Medina et al. 2014). We compared the changes in NET-seq signals as a function of these ratings, finding that higher sensitivity is strongly correlated with larger decreases in Pol II occupancy in *xrn*1Δ, *dhh*1Δ, and *lsm*1Δ; a weaker pattern was apparent for *ccr*4Δ and none for *rpb*4Δ (Fig. 1C). Our findings directly support earlier classification of genes into the Xrn1 synthegradon using GRO (Medina et al. 2014) and additionally demonstrate that transcription of the same genes is also activated by Dhh1, Lsm1, and Ccr4. Rpb4’s role in transcription is unrelated to this classification, most likely because it affects the transcription of most, if not all, genes (Schulz et al. 2014).

To understand whether particular classes of genes are more affected by DF deletions, we ranked genes according to their fold changes in total Pol II occupancies. Genes whose Pol II counts decreased or increased significantly were called “up-regulated” (normally their transcription is induced by the concerned DFs) or “downregulated” genes (normally their transcription is repressed by the concerned DFs), respectively. Briefly, we found that the most affected genes in *xrn*1Δ strains are those which are required for cell proliferation under optimal conditions when glycolysis is the main producer of ATP (and aerobic metabolism is partially repressed). For example, deletion of *XRN*1 results in reduced transcription of ribosomal protein (RP), ribosome biogenesis (RiBi), and transcription factor (TF) genes (Fig. S4A) and increased transcription of aerobic metabolic genes (cellular respiration, mitochondria, ATP synthesis and transport, and cytochromes) (Fig. S4B). Highly similar classes of affected genes were identified among Dhh1-upregulated genes in *dhh*1Δ (results not shown). Since the deletion of *XRN*1 results in lower Pol II levels in RP and RiBi genes but increased levels in aerobic metabolic genes, it seems that Xrn1 is involved in the balance between building cell mass and the metabolism. Interestingly, Xrn1 is regulated by Snf1-activated phosphorylation (Braun and Young 2014) and *XRN*1 interacts genetically with *T OR*2 (Costanzo et al. 2016). Snf1 and Tor2 are kinases that function in a similar balance between building cell mass and metabolism.

### Deletions of mRNA decay factors affect Pol II occupancy in both ends of transcription units of protein coding genes

A notable feature of NET-seq is that it can capture arrested Pol II in addition to those which are productively elongating. While this may complicate direct estimation of transcription rates, it allows for a more refined interrogation of changes in Pol II processivity. For instance, the role of the elongation factor TFIIS (Dst1) in facilitating the release of backtracked Pol II was studied using NET-seq (Churchman and Weissman 2011). In the same vein, we examined whether our deleted genes affect Pol II distributions across genes by constructing metagene densities (see Methods). We first observed that WT samples displayed a ramp-like accumulation of reads ∼100 bp downstream of transcription start sites (TSS), in agreement with previous results (Churchman and Weissman 2011). Remarkably, TSS-proximal densities decreased strongly in *xrn*1Δ and *dhh*1Δ. In contrast, *ccr*4Δ and *rpb*4Δ exhibited even sharper Pol II occupancy profiles in these regions which also resided closer to TSS than those present in WT samples (Fig. 2). Pol II additionally pauses at the sites where Pol II transcripts are cleaved and post-transcriptionally polyadenylated, henceforth denoted polyadenylation sites (PAS) (Bentley 2014; Hyman and Moore 1993; Kazerouninia et al. 2010; Kuehner et al. 2011; Larson et al. 2011; Mischo and Proudfoot 2013). We observed a trend of higher densities near PAS across all mutants, with a particularly pronounced increase in *xrn*1Δ and *dhh*1Δ and smaller changes apparent in the remaining mutants (Fig. 2). Together, these results indicate that deletions of *XRN*1 and other DFs contribute to the high Pol II occupancy ∼100 bp downstream of TSS. We note that the 5’ and 3’ changes in Pol II occupancy are seemingly unrelated to growth rate given the lack of correlation between these changes (Fig. 2) and growth rates (Fig. S5). For example, both *xrn*1Δ and *rpb*4Δ cells grow more slowly than WT but display quite different metagenes. Likewise, *lsm*1Δ, *dhh*1Δ, and *ccr*4Δ grow similarly but present notably distinct average Pol II profiles. Conversely, *ccr*4Δ and *rpb*4Δ cells grow at different rates, yet possess similar metagenes.

**Fig. 2.**
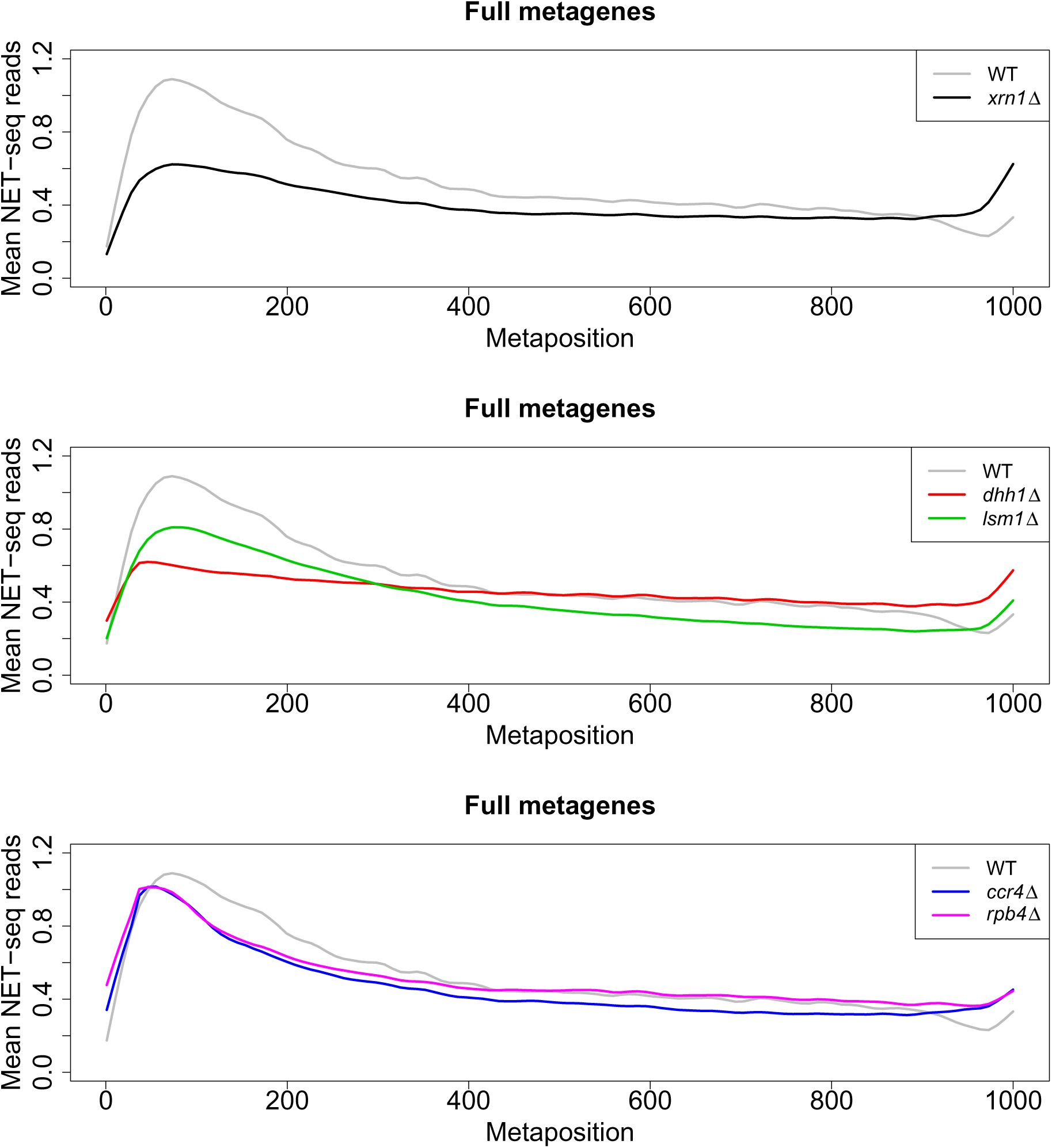
Comparison of normalized full-body metagenes. Normalized reads were aggregated within between the TSS and PAS. Genes were then re-scaled to each be of length 1000nt. Finally, the read counts corresponding to the new “metapositions” were averaged to yield a picture of Pol II occupancy along whole gene bodies. Different panels show comparisons between WT and the indicated deletion strains.

### Pol II initiation and elongation at the 5’ ends

Given the locations of the most apparent changes, we focused on the respective ends of transcription units, beginning with the 5’ end. To achieve finer resolution in this region, we generated metagene profiles for reads adjacent to TSS (Figs. 3A and S6). Consistent with our prior metagene analysis, we observed disruptions in normal promoter-proximal accumulation of NET-seq reads in *xrn*1Δ as well as *dhh*1Δ and *lsm*1Δ strains (Figs. 3A, S6A, S6B). Based on mathematical modeling (see below), we interpret these results to mean that the deletion of any one of these mRNA decay factors results in defective transcription initiation in addition to previous findings that *XRN*1 deletion leads to defective elongation (Haimovich et al. 2013; Begley et al. 2019). In contrast, deletions of *CCR*4 and *RPB*4 resulted in sharper Pol II peaks. In these two strains, Pol II occupancy was slightly closer to TSS than in the WT (Figs. S6C, S6D).

**Fig. 3.**
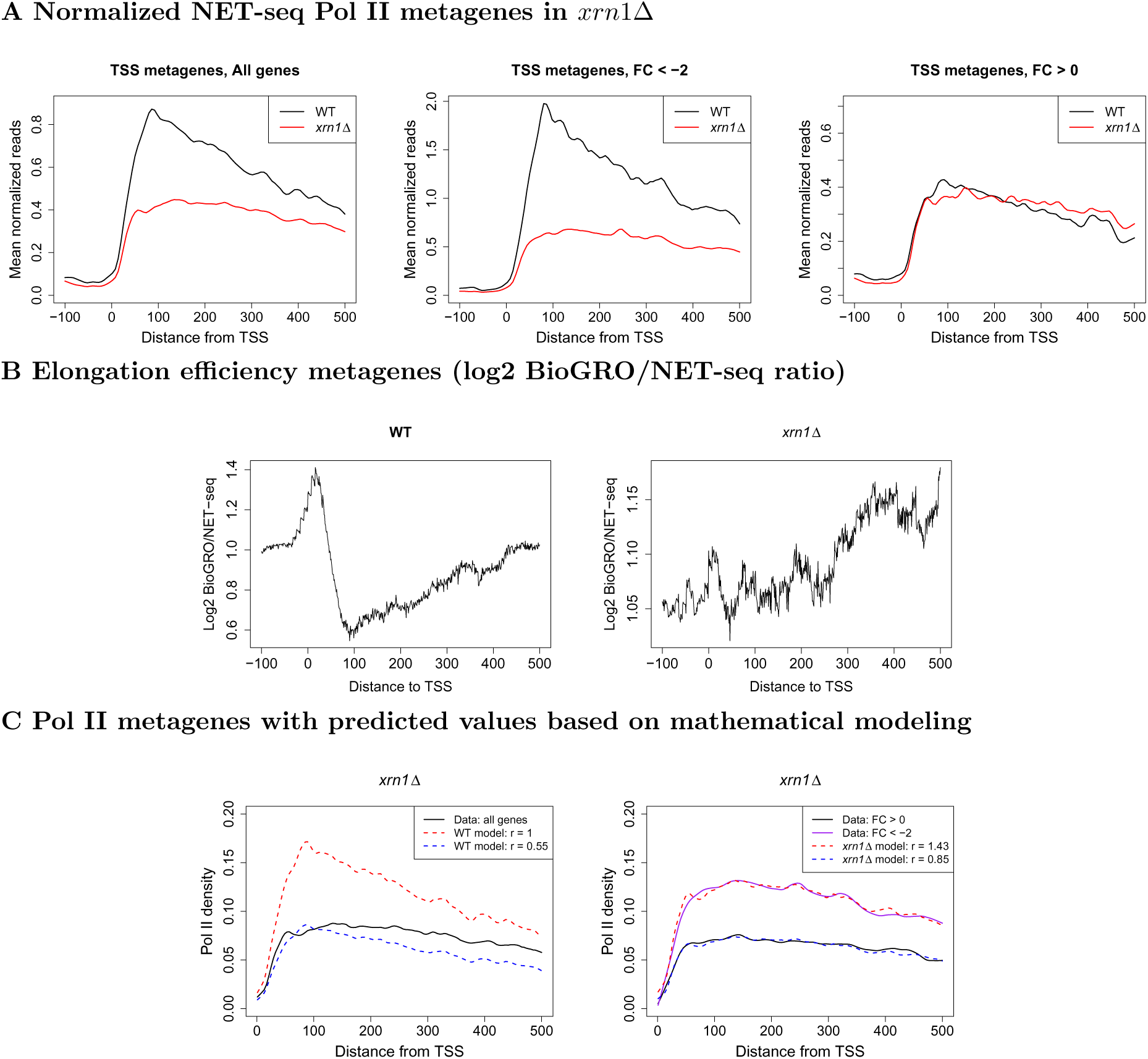
Metagene profiles near TSS in WT and *xrn*1Δ for Pol II and BioGRO/NET-seq ratios. (A) NET-seq reads were extracted (-100:500 relative to TSS), normalized, and averaged. Genes were separated into those which are stimulated (FC < 2) or repressed by Xrn1 (FC > 0). (B) We extracted BioGRO and NET-seq values in the -100:500 region with respect to the TSS for all genes. For each gene, we smoothed the BioGRO and NET-seq profiles and took the log2 of their ratios. We then averaged over all genes to yield elongation efficiency metagenes. (C) We applied a mathematical model (see Methods) to investigate how initiation and elongation rates affect metagenes. Elongation rates for WT and mutant metagenes were estimated and initiation rates (“r”) were varied to find the best fits. L - Varying initiation rates while using only the estimated WT elongation rates; R-varying initiation rates while using the estimated elongation rates from the *xrn*1Δ metagene. See Fig. S6 for other mutants.

We computed additional Pol II metagenes after stratifying genes into those which were up- or downregulated by Xrn1, finding that these two classes of genes responded fundamentally differently to the deletion of *XRN*1 (Fig. 3A). In genes strongly upregulated by Xrn1 (FC < −2), 5’ Pol II occupancy underwent a notable reduction in *xrn*1Δ strains. Differences between these gene classes were also apparent in WT cells; genes upregulated by Xrn1 are highly transcribed and exhibited relatively higher Pol II levels with steeper slopes in 5’ regions. In contrast, those which are downregulated did not (Fig. 3A), suggesting that Xrn1 deletion differentially affects genes based on their normal transcriptional patterns. One potential caveat is that the genes upregulated by Xrn1 have roughly three times as many reads as those it downregulated, so the observed differences may be related to the extent of transcription. Nevertheless, it is interesting that the presence of Xrn1 stimulates highly transcribed genes and represses lowly transcribed genes, helping maintain the gap in transcription levels. Consequently, upon disruption of Xrn1, we expect the genome-wide spread of transcription rates to shrink. Up- and downregulated genes were subsequently determined for the other deletion strains using the same criteria (Fig. S6). The resulting profiles from *lsm*1Δ and *dhh*1Δ cells were very similar to those of *xrn*1Δ, suggesting that Xrn1, Lsm1 and Dhh1 function similarly (Figs. 3A, S6A, S6B). On the other hand, changes in *ccr*4Δ and *rpb*4Δ profiles exhibited different patterns (Figs. S6C, S6D). upregulated genes displayed reductions in both mutants whereas 5’ peaks in downregulated genes increased and exceeded levels in the WT. We further computed Pol II metagenes upon restriction to specific classes of genes. Two examples were genes annotated to the ribosome biogenesis (RiBi) and ribosomal protein (RP) ontologies, which we selected due to their high expression levels and the reported effect of Xrn1 on their expression (Medina et al. 2014). TSS metagenes showed reductions in Pol II in both classes, although RP genes displayed heightened and/or sharper Pol II distributions in *ccr*4Δ, *dhh*1Δ, and *rpb*4Δ (Figs. S8A, S9A).

We sought to bolster our hypothesis of reduced elongation rates in *xrn*1Δ by looking for signs of increased pausing or backtracking. As backtracked Pol II cannot elongate because the nascent RNA is displaced from the active site (Churchman and Weissman 2011), they can be detected by NET-seq but not GRO because the latter assay relies on transcription elongation. Consequently, backtracking rates can be evaluated by comparing data from these two assays. To investigate transcriptional activity per unit Pol II - the elongation efficiency - in WT and *xrn*1Δ strains, we compared BioGRO data (Jordán-Pla et al. 2014; Jordán-Pla et al. 2016) and our NET-seq data. This is analogous to the analysis in Fig. S1B but with spatial resolution of Pol II activity. We focused on the two regions that demonstrated strong responses to Xrn1 deletion - the 5’ and 3’ ends, the latter of which is discussed in the subsequent section. In WT cells, we observed high elongation efficiency extending from TSS until ∼30 bp post-TSS, followed by a gradual decrease until ∼100 bp post-TSS (Fig. 3B). We thus propose that WT Pol II backtracks and pauses more often as it approaches ∼100 bp past-TSS. In cells lacking Xrn1, this initial high elongation efficiency region vanishes, suggesting dysregulation of these processes (Fig. 3B).

The accumulation of NET-seq reads at ∼100 bp downstream of TSS could represent a controlled Pol II pausing phenomenon akin to what has been described for many metazoan genes (see Introduction). Alternatively, the trademark buildup of Pol II near TSS may simply be the result of unbalanced initiation and 5’ elongation rates. To investigate the plausibility of the latter scenario, we employed a recently developed mathematical model which considers particles moving along a 1-dimensional path that was recently applied to ribosomes (Erdmann-Pham et al. 2018). We first used our computed metagene profiles to estimate reference initiation and site-specific elongation rates and then examined the results of perturbing these parameters. This allowed us to perform *in silico* experiments to separate the contributions of initiation and elongation rate changes and to infer the contributions of the studied DFs to elongation dynamics. Although Monte Carlo models of transcription have been considered in a handful of prior studies (Darzacq et al. 2007; Ehrensberger et al. 2013; Grosso et al. 2012; Jonkers et al. 2014; Le Martelot et al. 2012), to the best of our knowledge this is the first attempt to apply a model which rigorously and flexibly handles both spatial heterogeneity in elongation rates and the mutual interference of co-localized Pol II. Furthermore, it permits us to obtain analytical solutions from input parameters, increasing the precision of our analysis. Based on this model, we found that the observed WT Pol II metagene was indeed consistent with slower 5’ elongation compared to initiation (Fig. 3C). Thus, a controlled pausing event is not necessary to reproduce the observed profiles. Of course, our simulation does not conclusively rule out such a possibility; nevertheless, given the absence of supporting data in this work or the wider literature, we propose that imbalanced rates of transcription initiation and elongation constitute the major cause of 5’ Pol II accumulation in WT cells. Given the computed transcription elongation efficiency profiles (Fig. 3B), we additionally propose that the gradual decrease in Pol II processivity as Pol II approaches the 100 bp position exacerbates the imbalance between rates of initiation and elongation. Thus, the characteristic peak at ∼100 bp seems to result from the balance between initiation rates and position-dependent kinetics of Pol II elongation.

We next performed a similar analysis on mutant metagenes. To test whether the changes between mutant and WT profiles could be replicated by solely modulating initiation, we fixed elongation rates to the values inferred for the WT (see Methods) and varied initiation rates over a range of values in the simulation model. This procedure produced similar simulated profiles to those observed for *xrn*1Δ and *lsm*1Δ, suggesting that the deletion of these genes compromised transcription initiation. However, the observed NET-seq profiles were notably flatter than the simulated ones (Figs. 3C, S6B), indicating that defects in elongation in the mutants should also be considered. For the *ccr*4Δ, *rpb*4Δ, and *dhh*1Δ mutants, simulated profiles using WT elongation rates were unable to recapitulate the appearance of pronounced peaks slightly upstream of 100 bp (Figs. S6A, S6C, S6D).This indicates that, for these strains, our observed metagene profiles cannot be explained by simple changes in the overall balance between initiation and elongation. Hence it is likely that more complex kinetics are involved in which the ∼ 100 bp location may serve as a transition point. This notion is supported by the clear differences in behavior observed at the 30 bp and 100 bp positions post-TSS in the WT and *xrn*1Δ elongation efficiency (BioGRO/NET-seq) profiles (Fig. 3B). In summary, while the differences between heights of 5’ peaks in WT and mutant strains can be explained by reduced initiation rates, the differences in profile shapes cannot be totally accounted for by manipulating this single quantity. Hence transcription elongation is affected both before and after the 100 bp mark in *xrn*1Δ cells (Fig. 3). We therefore propose that initiation rate reduction is a major consequence of *XRN*1 and *LSM*1 deletion with additional decreases also occurring in elongation rates. Furthermore, although the respective deletions of *DHH*1, *CCR*4, and *RPB*4 also reduce initiation rates, they have additional targeted effects on elongation rates in the first 100 bp of genes which differ from those of Xrn1 and Lsm1.

### Deletion of mRNA decay factors leads to a marked accumulation of Pol II near PAS, probably due to increased pausing/backtracking

WT Pol II pauses at PAS (Fig. 4A), probably to provide time for the PA mechanism to function (Tian and Manley 2017). In the *xrn*1Δ, *dhh*1Δ, and *lsm*1Δ strains, abnormally high spikes were observed in this region (Figs. 4A, S7). This pattern suggests enhanced pausing in the absence of these DFs. Interestingly, mutant strains displayed abnormally high accumulations of reads beginning ∼75 bp upstream of PAS and lasting until PAS. Downstream of these PAS, NET-seq reads accumulated due to transcription that continues beyond PAS before reaching transcription termination sites (Bentley 2014). Atypically low NET-seq reads were observed downstream of PAS in the *xrn*1Δ mutant strains, suggesting that less Pol II could be released from a paused state in the absence of Xrn1. In the *rpb*4Δ and *ccr*4Δ strains, we detected accumulations of reads upstream and downstream of PAS, but the actual PAS peaks were comparable to those in the WT (Fig. S7). Separation into genes up- and downregulated by Xrn1 revealed 3’ occupancy patterns unlike those in 5’ ends. Whereas upregulated genes (FC < −2) had displayed large reductions in 5’ Pol II levels (Fig. 3A), 3’ occupancy demonstrated relatively little sensitivity to the presence of Xrn1 (Fig. 4A). Genes downregulated by Xrn1 (FC > 0) displayed nearly the opposite behavior, as 5’ occupancy was insensitive to *XRN*1 deletion while 3’ pausing greatly increased (Figs. 3A, 4A). Deletions of *CCR*4 and *RPB*4 had smaller effects on pausing at PAS, although general occupancy increases were present in downregulated genes (Fig. S7).

**Fig. 4.**
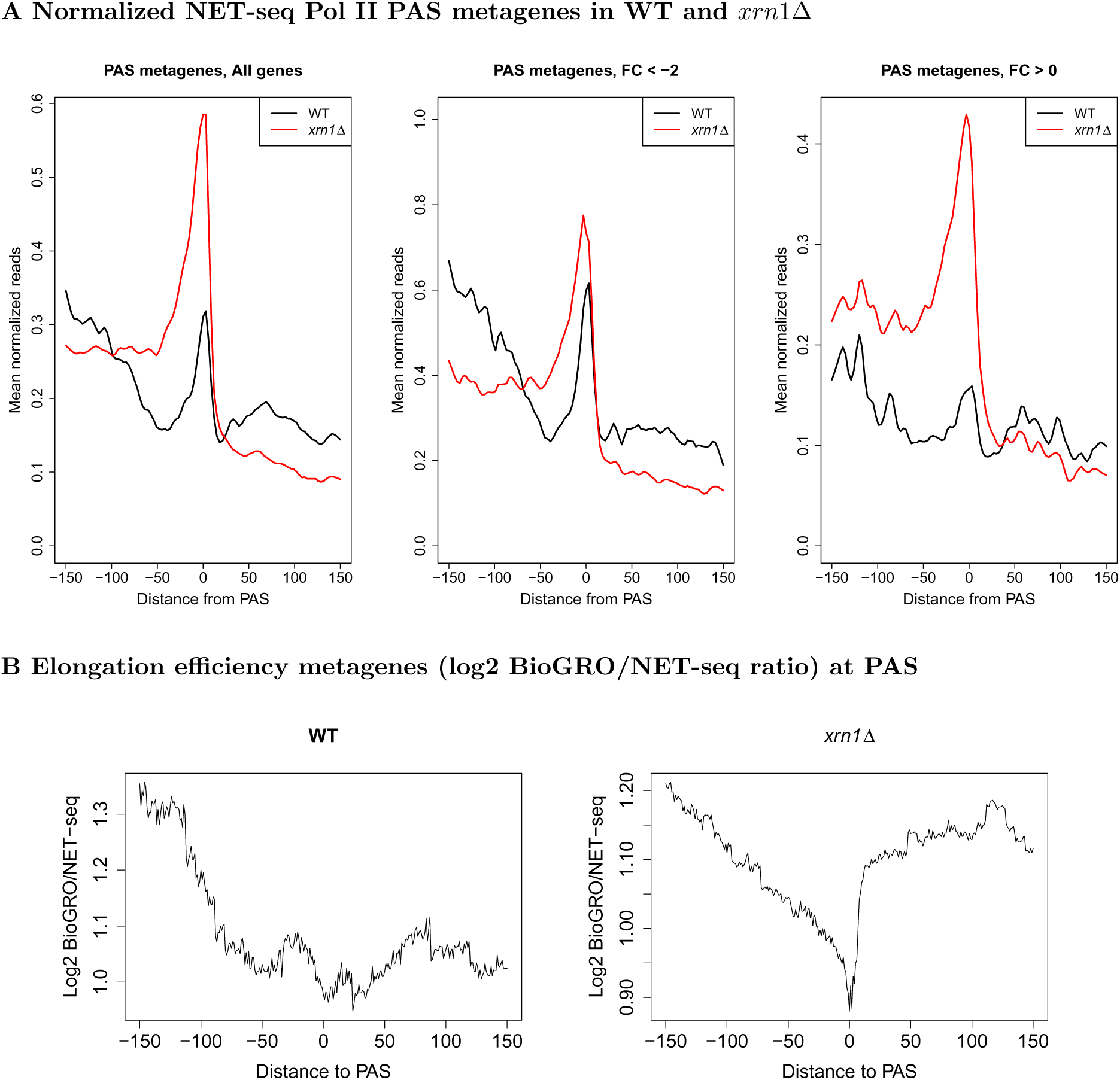
Metagene profiles near PAS in WT and *xrn*1Δ for Pol II and BioGRO/NET-seq ratios. (A) We extracted NET-seq reads (-150:150 relative to PAS), normalized, and averaged. Genes were separated into those which are stimulated (FC < 2) or repressed by Xrn1 (FC > 0). (C) We extracted BioGRO and NET-seq values in the -150:150 region with respect to the PAS for all genes. For each gene, we smoothed the BioGRO and NET-seq profiles and took the log2 of their ratios. We then averaged over all genes to yield elongation efficiency metagenes.

Our metagene analyses showed that *xrn*1Δ and *dhh*1Δ cells accumulate abnormally small numbers of reads in 5’ regions and unusually high numbers of reads in 3’ regions (∼-100 until PAS). Because these analyses aggregated reads across all genes, it was unclear whether profile changes were driven by widespread behavior or simply a small number of highly impacted genes. Given that we observed somewhat similar 5’ and 3’ profiles in RiBi and RP genes (Figs. S8, S9) as well as SAGA and TFIID-dominated genes (Figs. S10, S11), we suspected that many genes were affected. To address this issue more directly, we determined Pol II 5’/3’ ratios for each mutant, an approach which preserves the pairing of 5’ and 3’ occupancies in individual genes (Fig. S12). Indeed, Pol II 5’/3’ ratios were substantially lower in *xrn*1Δ compared to WT strains, and the overall shift in the distribution of these ratios suggests that the changes in the 5’ and 3’ metagene profiles are not confined to a small number of genes. Interestingly, the *dhh*1Δ strain exhibited a similar pattern of Pol II 5’/3’ ratios, and additional modest decreases in Pol II 5’/3’ ratios were observed in the remaining mutants.

Much like before, we generated BioGRO/NET-seq ratio profiles to consider the role of pausing and backtracking near PAS (Fig. 4B). We found that WT Pol II elongation efficiency gradually decreased as it moved towards PAS while *xrn*1Δ cells displayed a precipitous efficiency reduction at the PAS. Thus, Pol II that accumulate upstream of PAS in *xrn*1Δ cells are relatively inactive, both in WT but notably more so in *xrn*1Δ cells, most likely in a backtracked configuration. Collectively, the most pronounced effects of DFs in transcription are at the ends of genes where Pol II processivity decreases in the mutant strains, potentially relating to increased Pol II backtracking. Moreover, the differences between DF deletion-induced responses indicate additional defects in transcription initiation. These differences cannot be simply attributed to growth rates because *lsm*1Δ cells proliferate comparably to *dhh*1Δ and *ccr*4Δ (Fig. S5) despite different effects of the deletions of *LSM*1, *DHH*1, and *CCR*4 on metagene profiles (Fig. S6).

### Transcription termination (beyond PAS) seems to be affected by deletion of the studied mRNA decay factors

Transcription termination, which occurs downstream of PAS, is allosterically modulated by the PA mechanism (Richard and Manley 2009; Tian and Manley 2017). Because we found that our studied DFs function in the PA process, we examined whether transcription termination is also affected by the deletion of DFs. Direct analysis of changes in termination using NET-seq is challenging because it does not identify transcription termination efficiently, probably because there are multiple termination events (Churchman and Weissman 2012). Thefore, we examined the effect of DFs on this process indirectly by taking advantage of the capacity of ChIP-seq, or NET-seq in our case, to report Pol II pausing due to collisions of two convergently transcribed Pol II molecules (Hobson et al. 2012). After defining the midpoint between convergent genes as the halfway point between the ends of paired 3’ UTRs, we found that the NET-seq signal in WT cells decreases gradually as a function of distance from the 3’ ends, consistent with gradual termination post-PAS. In contrast, mutant strains displayed accumulations of Pol II near the midpoints of convergent gene pairs as evidenced by midpoint peaks (Fig. 5). Our results are reminiscent of the previous demonstration of the Pol II buildup between convergent genes in strains lacking Elc1, a protein which aids in the removal of stalled Pol II (Hobson et al. 2012). This raises an alternative explanation in which the studied DFs stimulate the degradation of colliding Pol II. To verify that the accumulation between convergent gene pairs was truly a result of Pol II collisions, we stratified genes based on the distances between their respective PAS. This analysis demonstrated that Pol II occupancy between such pairs gradually decreased as a function of the distance from the midpoints (Fig. S13), suggesting that as the distance between respective PAS increases, Pol II has more opportunities to terminate in both WT and mutant strains. This is consistent with a model wherein Pol II normally terminates within a window 100-200 bp downstream of PAS but instead continues to transcribe further downstream in mutant strains because of less efficient termination. As a point of reference, we also performed this analysis for divergent pairs, finding only the expected differences due to reduced Pol II occupancy downstream of TSS (Fig. 5B).

**Fig. 5.**
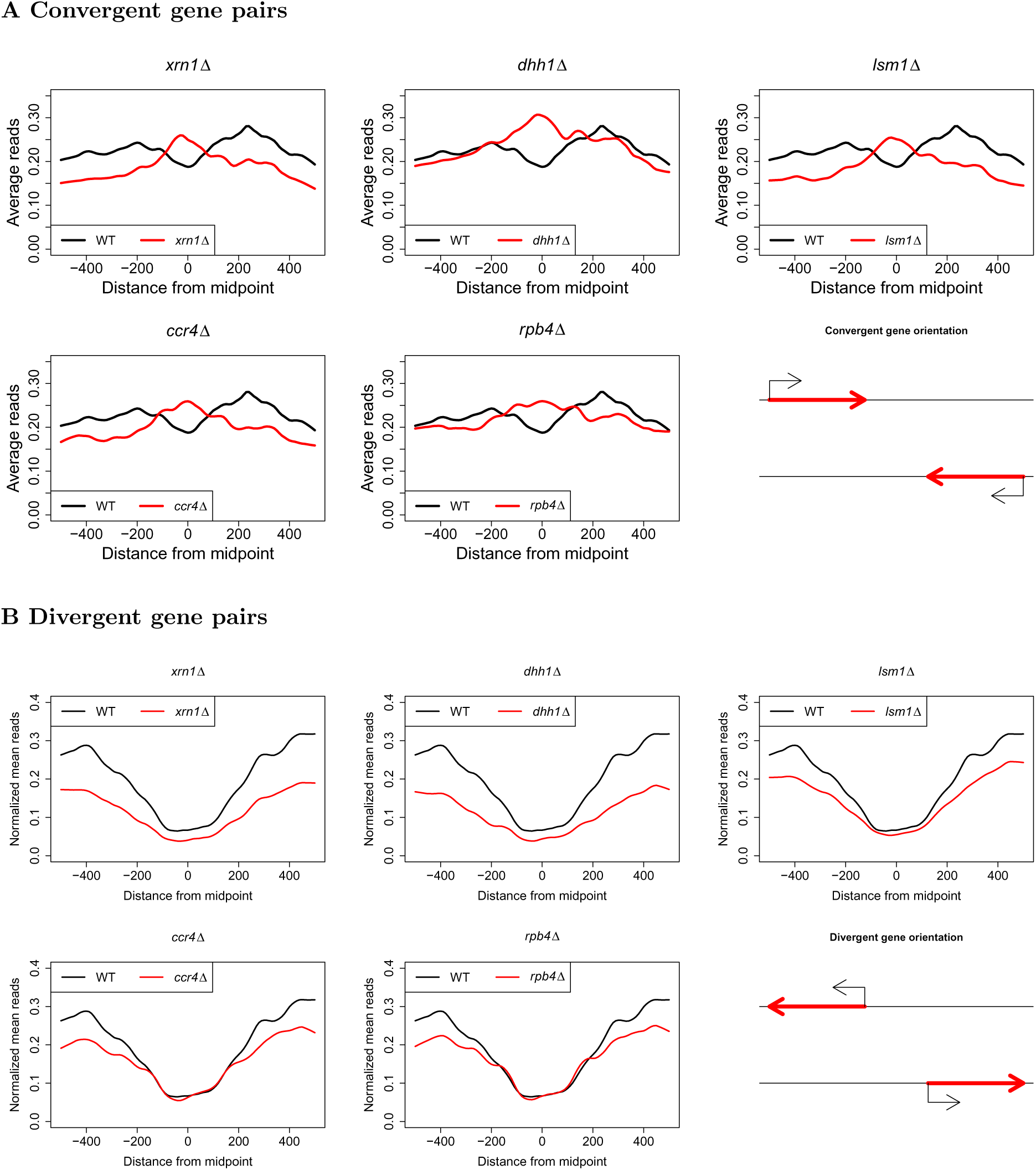
Metagenes for convergent and divergent gene pairs. Convergent and divergent gene pairs were determined by the lengths between their PAS (convergent) and TSS (divergent). Midpoints between genes were defined as the halfway point between these respective features, and gene distances were computed as the difference between the annotated features on the negative and positive strand, respectively. Normalized NET-seq reads were then extracted for sites within 500 bp of gene midpoints and subsequently averaged to produce the metagene profiles.

### Transcription in non-coding regions is also compromised by DF deletions

NET-seq provides an opportunity to monitor the production of unstable transcripts because it quantifies bound Pol II rather than mature RNAs and is little affected by RNA stability. Consequently, we investigated the effect of DF deletions on the transcription of non-coding RNAs (ncRNAs) by considering the changes in Pol II occupancy at chromosomal loci encoding cryptic unstable transcripts (CUTs), Nrd1-unterminated transcripts (NUTs), stable unannotated transcripts (SUTs), and Xrn1-sensitive unstable transcripts (XUTs). Both CUTs and SUTs frequently originate in the nucleosome free regions (NFRs) upstream of sense promoters and often run antisense to protein-coding genes, posing the possibility of *cis*-regulatory roles (Bumgarner et al. 2009; Tisseur et al. 2011). Most CUTs are rapidly degraded by the exosome while SUT degradation is more reliant on Xrn1 activity (Xu et al. 2009), though it has been noted that some CUTs are also degraded by Xrn1 (Marquardt et al. 2011). XUTs comprise an additional class of regulatory ncRNAs which are degraded by Xrn1 and whose transcript levels increase substantially in Xrn1’s absence (van Dijk et al. 2011). As CUTs and SUTs are to varying degrees degraded by Xrn1 and XUTs are by definition sensitive to its deletion, nascent transcriptional changes in the *xrn*1Δ samples are especially relevant to understanding the interconnectedness of RNA synthesis and decay for non-coding transcripts. Finally, NUTs are transcripts whose termination is altered after the nuclear depletion of Nrd1, resulting in longer transcripts (Schulz et al. 2013). Computation of NET-seq fold changes across gene bodies demonstrated global reductions in Pol II occupancy in *xrn*1Δ, *dhh*1Δ, and *rpb*4Δ (Fig. S14). Interestingly, the effect of deleting *XRN*1, *DHH*1, and *RPB*4 on the transcription of CUTs and NUTs was higher than their effect on the transcription of coding genes. In contrast, no effect of deleting *LSM*1 on ncRNA transcription was observed. Somewhat unexpectedly, Pol II occupancy in NUT loci was highly affected by the deletion of *XRN*1 whereas that of XUTs, degraded mainly by Xrn1 (van Dijk et al. 2011), showed the smallest sensitivity to *XRN*1 deletion (Fig. S14).

To see if distributional changes in Pol II also appeared in ncRNAs, we again computed meta-gene densities (Fig. S15). The impact of DFs at TSS of all ncRNA types resembled those of protein-coding genes. Specifically, deletion of *RPB*4 or *CCR*4 had little effect, whereas deletion of other DFs decreased 5’-proximal Pol II occupancy. Interestingly, however, for none of the metagenes did we observe the increase in Pol II adjacent to PAS that was characteristic of protein-coding genes, suggesting that the deleted proteins interact differently with the PA machinery of coding and non-coding genes. Together, these results may signify that the considered DFs regulate initiation or early elongation in ncRNAs but have no impact on termination, perhaps as a consequence of differences in the respective mechanisms of polyadenylation in coding genes and genes encoding ncRNAs.

Despite the strong effect of deleting some DFs on transcription of ncRNAs, we found small anticorrelations between sense and antisense Pol II occupancy, both in WT and deletion strains (Fig. S16A). Unlike convergent pairs, transcription levels in divergent pairs showed moderate positive correlations. Broadly speaking, correlations between sense and antisense transcription are roughly equal no matter if the antisense transcript is a coding gene or a ncRNA. Thus, there seems to be no obvious global relationship between coding and ncRNA genes, consistent with what has been reported previously (Murray et al. 2015). Reduced transcription levels in genes with overlapping convergent transcripts implies that the weak anticorrelation that found among convergent pairs is primarily driven by the presence or absence of antisense transcripts rather than the transcription levels thereof (data not shown), in agreement with previous proposals (Wery et al. 2018). It is possible that the positive correlations among divergent pairs were simply due to the common chromatin environment of nearby divergent promoters (Murray et al. 2015; Xu et al. 2009). In summary, we could not find any indication that DFs target ncRNA genes in order to modulate transcription in protein-coding genes. In fact, we found that genes which are highly sensitive to *XRN*1 deletion (i.e. the Xrn1 synthegradon) are less likely to have convergent antisense transcripts of any type (results not shown).

### Transcriptional responses to DF deletions differ between SAGA- and TFIID-dominated genes

Prior studies have classified genes as SAGA- or TFIID-dominated according to the measured changes in mRNA levels after inactivation of central components of the SAGA (mainly Spt3) and TFIID (mainly Taf1) complexes (Basehoar et al. 2004; Huisinga and Pugh 2004). Genes which were classified as “SAGA-dominated” comprise about 10% of the yeast genome, many of which are highly responsive to the environment and are likely to have TATA boxes in their core promoters (Huisinga and Pugh 2004). Meanwhile, “TFIID-dominated” genes make up most of the remaining 90% of genes and are frequently housekeeping genes and containing TATA-like elements in their core promoters (Huisinga and Pugh 2004). However, analyses such as these which interrogate changes in mRNA levels are unable to distinguish between the contributions of mRNA synthesis and decay. Recently, it was discovered that all promoters recruit both SAGA and TFIID (Baptista et al. 2017; Bonnet et al. 2014; Warfield et al. 2017); mutations in either complex resulted in defective transcription, but in most cases mRNA levels were unaffected due to feedback mechanisms (Baptista et al. 2017; Bonnet et al. 2014; Warfield et al. 2017). We therefore examined whether our studied DFs are differentially involved with the two groups of genes by comparing the fold changes in Pol II levels of genes in each group after deletions of DFs (Fig. 6A). As expected, we found decreases in Pol II occupancy in TFIID genes due to *XRN*1 deletion. However, median Pol II occupancy in “SAGA-dominated” genes remained unaffected in *xrn*1Δ, *lsm*1Δ, and *rpb*4Δ strains and even slightly increased in *ccr*4Δ and *dhh*1Δ strains (Fig. 6A). Note, however, that these results refer to Pol II occupancy along the entire gene. We next compared fold changes in NET-seq reads near TSS and PAS (Fig. 6B), finding that DF deletions affected TFIID-dominated genes more than SAGA-dominated genes.

**Fig. 6.**
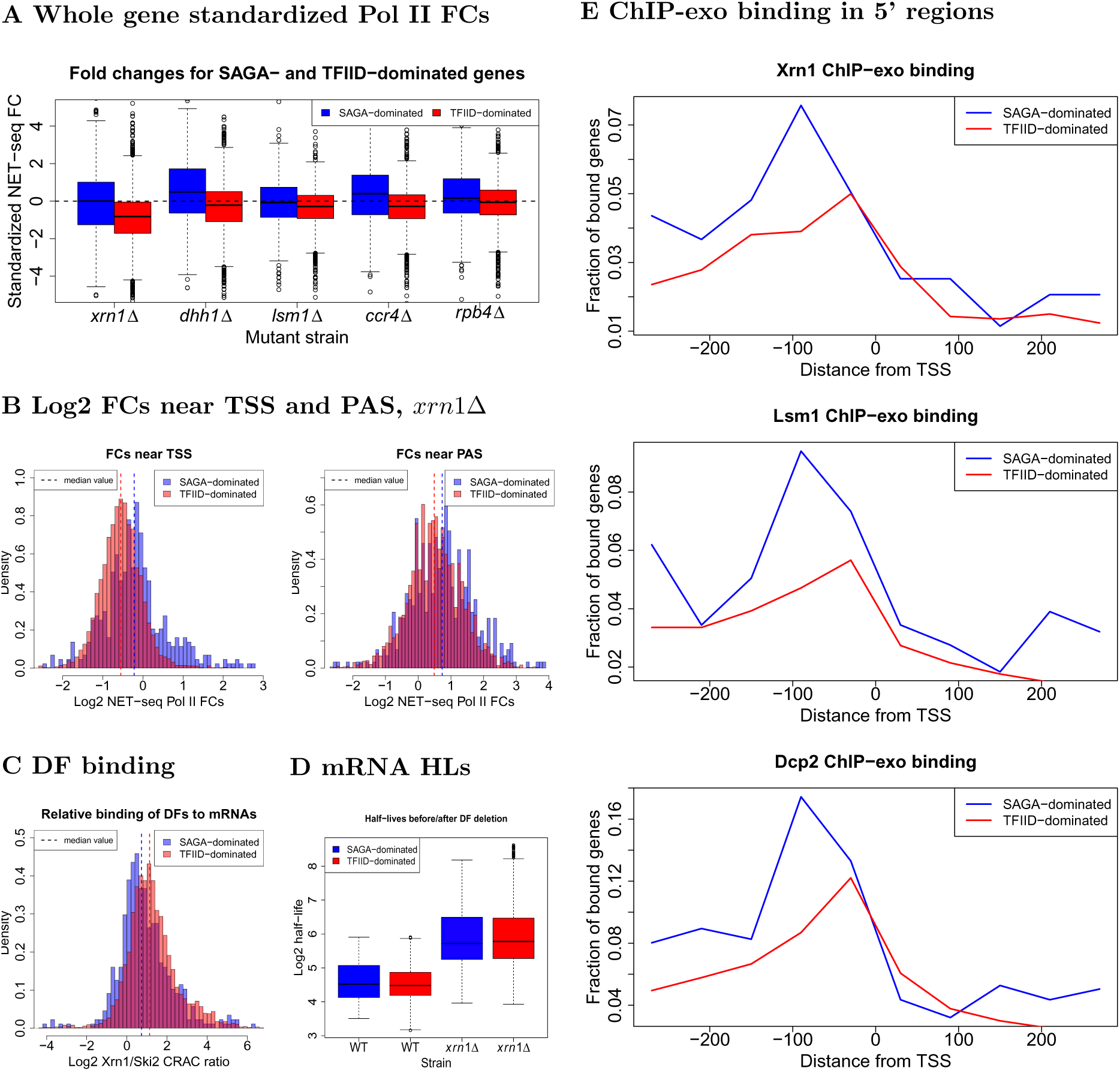
Comparison of transcription, decay, and protein binding for SAGA- and TFIID-dominated genes. (A) Fold changes, computed as in Fig. 1A, for SAGA- or TFIID-dominated genes, as indicated. (B) Histograms of log2 NET-seq Pol II FCs for regions near TSS (-100:500) and PAS (-150:150) in *xrn*1Δ. (C) Xrn1 and Ski2 CRAC data (Tuck and Tollervey 2013) mapped to each gene were summed and the log2 ratio of mapped reads for each DF in each gene was taken. (D) Comparison of mRNA half-lives before and after XRN1 deletion (Medina et al. 2014). (E) Reads were binned into windows of 60 bp starting 300 bp upstream and extending 300 bp downstream of TSS. The proportion of bins having more than 10 recorded reads was then computed across the genome and plotted (Haimovich et al. 2013).

The discrepancy between mRNA levels and transcription rates suggests that an important aspect of the division between SAGA- and TFIID-dominated genes may lie in the regulation of transcript decay, possibly mediated by DFs (see above). To explore this possibility, we analyzed publicly available UV cross-linking and analysis of cDNA (CRAC) data for Ski2 and Xrn1 (Tuck and Tollervey 2013). Whereas Xrn1 is the key enzyme of the 5′ − 3′ mRNA decay process, Ski2 serves as a vital component of the exosome and thus represents the 3′ − 5′ cytoplasmic decay pathway. We found that each of these proteins bound to both classes of genes, suggesting that they play a role in the decay of transcripts regardless of complex annotations (Fig. 6C). However, both Ski2 and Xrn1 bind more frequently to SAGA-dominated genes than TFIID-dominated genes even after accounting for transcript length and steady-state mRNA levels. The discrepancy was stronger for Ski2, as this protein bound to mRNAs from SAGA-dominated genes at nearly twice the rate compared to TFIID-dominated genes (∼ 1.86×). Moreover, we estimated Xrn1 binding for SAGA-dominated transcripts to be roughly 1.47× the rate as for TFIID-dominated transcripts. That binding of Ski2 occurs at a higher comparative rate between classes than Xrn1 suggests that the decay of SAGA-dominated transcripts is more dependent on the exosome than the Xrn1-led 5′ − 3′ pathway. At any rate, SAGA- and TFIID-dominated genes are differentiated by Xrn1 and Ski2 binding. However, despite the higher binding rates for both proteins, a comparison of the half-lives of mRNAs from SAGA- and TFIID-dominated genes based on previously published data (Medina et al. 2014) failed to detect a discernible difference (Fig. 6D). Thus, on one hand, both the SAGA and TFIID complexes regulate transcription of all genes (Baptista et al. 2017; Bonnet et al. 2014; Warfield et al. 2017) and their respective transcripts have comparable half-lives (Fig. 6D); conversely, transcription of SAGA- and TFIID-dominated genes is differentially affected by deletions of the studied DFs (Figs. 6A, 6B) and binding of Xrn1 and Ski2 to their respective transcripts differentiates between SAGA- and TFIID-dominated genes (Fig. 6C).

Inspired by the different effects of DFs on SAGA and TFIID genes, we explored the DNA binding patterns of three decay factors by analyzing previously generated ChIP-exo data for Dcp2, Lsm1, and Xrn1 (Haimovich et al. 2013) after stratifying genes according to their “classical” SAGA/TFIID labels. We found that all three proteins bind to SAGA-dominated genes at higher frequencies than TFIID-dominated genes (Fig. 6E). These proteins tend to bind further upstream of TSS for SAGA-dominated genes (peak ∼ 90 bp) compared to TFIID-dominated genes (∼ 30 bp), further highlighting the distinction between the two classes. Given the recent finding that SAGA localizes further upstream than TFIID and binds more frequently in SAGA-dominated than TFIID-dominated promoters (Baptista et al. 2017), it is possible that the interactions of Dcp2, Lsm1, and Xrn1 with promoters are influenced by the positions of bound SAGA and TFIID complexes as well as their binding frequencies. The higher binding rates of each protein to SAGA-dominated genes is somewhat surprising given that transcription of SAGA-dominated genes is less impacted by DF deletion, but perhaps reflects the observation that Xrn1 prefers to bind mRNAs of SAGA-dominated genes over those which are TFIID-dominated. It is also possible that the respective ChIP-exo profiles differ due to technical artifacts arising from the accessibility differences of the TAP-tag used to pull down DFs due to differing configurations of DFs within the two complexes. In any event, these results highlight the capacity of the studied DFs to differentiate between SAGA- and TFIID-dominated genes.

In summary, transcription in both SAGA- and TFIID-dominated genes is dependent on the SAGA and TFIID complexes, and their mRNA products have comparable half-lives in WT and in *xrn*1Δ strains. However, SAGA- and TFIID-dominated genes differ by (i) the effect that the studied DFs have on their transcription, (ii) chromatin binding features of Xrn1, Lsm1 and Dcp2, and (iii) the binding of Xrn1 and Ski2 to their mRNAs.

## Discussion

In recent years, interest in understanding the cross talk between mRNA synthesis and decay has grown. Under optimal proliferation conditions, various mRNA decay factors are involved in mRNA “buffering”, a feedback mechanism that minimizes changes in mRNA levels. In this coupling, reductions in either mRNA synthesis or decay are associated with compensatory reductions in the other process, resulting in relatively consistent concentrations of mRNAs. Although Xrn1 was identified as an effector of buffering, its mode of action has remained controversial (see Introduction). Using NET-seq, we found that the deletion of *XRN*1 generally resulted in the downregulation of transcription (Fig. 1A) and notably reduced the elongation efficiency of Pol II (BioGRO/NET-seq, Figs. 3B, 4B), consistent with a role for Xrn1 as a stimulator (Haimovich et al. 2013; Medina et al. 2014) rather than repressor (Sun et al. 2013) of transcription. As a transcriptional activator, Xrn1 primarily targets genes required for proliferation under optimal conditions, when cells are dependent mainly on fermentation (Figs. S4B, S4C). These GO terms are similar to those that characterize the Xrn1 synthegradon group identified by a GRO-based analysis (Medina et al. 2014). We also found that the absence of Xrn1 results in increased Pol II levels of a minor population of relatively lowly expressed genes that mainly encode proteins related to aerobic metabolism (Figs. S4A, S4C). Given earlier findings of direct binding of Xrn1 and Lsm1 (Haimovich et al. 2013) and our finding that these proteins bind promoters of both stimulated and repressed genes as well (results not shown), we suspect that this effect is direct. Thus, Xrn1 seems to be one of the factors that coordinates gene expression to permit efficient proliferation when fermentation is preferred. Our work also uncovered an underlying function by which Xrn1 targets transcription initiation and Pol II processivity, possibly via pausing and backtracking. Recent studies have reported that the release of promoter-proximal paused netazoan Pol II is a crucial component of the regulation of transcription under both optimal (Sheridan et al. 2019) and stress conditions (Bartman et al. 2019; Sheridan et al. 2019) (see Introduction). Though not as severe as in *S. pombe* or metazoans, promoter proximal Pol II accumulation has been reported in S. cerevisiae (Churchman and Weissman 2011; Baptista et al. 2017; Feldman and Peterson 2019), but the underlying cause is uncharacterized (Adelman and Lis 2012). This early build-up of Pol II has been implicated as a “checkpoint” of Pol II elongation regulated by the CTD kinase Kin28 (Rodríguez-Molina et al. 2016). Moreover, depletion of sirtuin proteins (Hst3 and Hst4) increases 5’ proximal accumulation (Feldman and Peterson 2019). Thus, although the correspondence to mammalian Pol II pausing remains unclear, it seems that Pol II is subject to promoter-proximal regulation in *S. cerevisiae* as well. Here we show that Xrn1 and other yeast DFs also likely modulate this type of regulation in genes which they upregulate. We also note reductions in initiation rates for many genes across mutants and demonstrate via mathematical modeling that both initiation and elongation rate reductions are necessary to recapitulate observed Pol II profiles. Furthermore, we showed that while initiation rate changes appear to be restricted to a subset of genes, elongation rate changes are more ubiquitous, as metagenes of both activated and repressed genes displayed flatter Pol II profiles in mutants than the WT (Figs. 3A, 3C, S6). Recently, Pol II was shown to frequently backtrack in promoter-proximal regions of human genes, with TFIIS-stimulated RNA cleavage helping to release Pol II from pause sites (Bartman et al. 2019; Sheridan et al. 2019). Pol II backtracking can be evaluated by comparing BioGRO to NET-seq because backtracked Pol II cannot elongate in vitro. Hence, whereas NET-seq putatively captures all bound Pol II, BioGRO only detects those which are productively elongating, and the elongation efficiency can be determined by considering the ratio of BioGRO to NET-seq reads. High elongation efficiency was observed in WT cells for the first ∼30 bp down-stream of TSS (roughly coincident with capping), followed by a gradual drop until around 100 bp downstream, suggesting that Pol II backtracks with increasing frequency as it approaches the ∼100 bp mark post-TSS, as was found in mammalian cells (Bartman et al. 2019; Sheridan et al. 2019). In the absence of Xrn1, this pattern is disrupted (Fig. 3B), suggesting that the normal regulation of backtracking is compromised. DFs have previously been implicated to function in the context of backtracking. For example, Ccr4 has been shown to physically interact with TFIIS and to directly modulate backtracking in conjunction with TFIIS (Dutta et al. 2015; Kruk et al. 2011). Likewise, Dhh1, Pat1, and Lsm1 also interact with TFIIS both physically and genetically (Collins et al. 2007; Costanzo et al. 2010; Costanzo et al. 2016; Dutta et al. 2015; Kuzmin et al. 2018; Miller et al. 2018; Srivas et al. 2016; Wilmes et al. 2008). Recently, Xrn1 was shown to increase TFIIS recruitment to Pol II (Begley et al. 2019), consistent with the role we assign to Xrn1 as a regulator of backtracking. All these observations provide outside credibility to our proposals. Why, then, do we detect differences in the impact of different DFs on 5’ Pol II accumulation? For example, whereas the deletion of *CCR*4 led to enhanced Pol II accumulation (Fig. S6C), the deletion of *XRN*1 led to the obliteration of Pol II accumulation at this position (Fig. 3A). The simplest explanation is that different DFs differentially affect the initiation/elongation ratio by targeting initiation, elongation/backtracking, or both as demonstrated using our mathematical model. This is consistent with the recent report that Xrn1 and Ccr4 differentially regulate Pol II elongation; namely, whereas deletion of *XRN*1 led to increased TFIIS-Pol II interaction, that of *CCR*4 had the opposite effect (Begley et al. 2019). Nonetheless, the exact mechanism remains to be determined. These results do not simply provide insight into a plausible mechanism by which Xrn1 affects transcription, as they also highlight the need to study pausing and backtracking in the first ∼100 bp of transcription units as possible regulatory steps in yeast transcription.

Deletions of *XRN*1 and other DFs also affects Pol II occupancy at PAS. Pausing at PAS in WT cells is more apparent in genes activated by DFs rather than those which are repressed, and deletion of *XRN*1, *DHH*1, or *LSM*1 led to notable increases in pausing at PAS of repressed genes. Additional perturbations manifest as abnormally high read accumulations in the regions beginning roughly 75 bp upstream of PAS and extending to PAS. We found that as Pol II approaches PAS its activity gradually decreases and that this reduction is intensified in *xrn*1Δ strains (Fig. 4B). One possibility is that Xrn1 sterically impedes backward movement of Pol II (i.e., backtracking), as was proposed for Ccr4 (Kruk et al. 2011; Dutta et al. 2015). Accordingly, in *xrn*1Δ and the other mutant strains, backward motion of Pol II is not repressed and therefore backtracking is enhanced, burdening the TFIIS-stimulated RNA cleavage process and leading to an accumulation of reads upstream of and at PAS. While our data highlight a unique feature of PAS and the region immediately upstream of them, more work is required to pinpoint the exact regulatory mechanisms governing transcription in these regions. It hence remains possible that unique chromatin architecture and DNA/RNA sequences combined with the recruitment of PA factors, TFIIS, and some DFs in this region are involved in regulating proper Pol II elongation rate and PA.

Transcription termination occurs downstream of PAS (Mischo and Proudfoot 2013). Cleavage and polyadenylation factors are presumed to act in the Pol II release step of transcription termination by allosterically modifying the properties of the transcription elongation complex (for a recent review, see Porrua et al. 2016). Our studied DFs, which are involved in cleavage and polyadenylation, further appear to affect termination as well. This was determined indirectly because NET-seq does not identify transcription termination efficiently (Churchman and Weissman 2012), probably because, for any single gene, termination does not occur at a single locus. Correspondingly, we found that the NET-seq signal decreases gradually as a function of distance from the 3’ ends, consistent with a gradual termination post-PAS. In contrast, the signals in several of the mutant strains, most notably *rpb*4Δ and *ccr*4Δ, increased relative to the WT. We interpreted these results to indicate that transcription termination post-PAS is less efficient in our mutant cells, thus increasing the probability that two opposing Pol II would collide.

We also studied the effects of DF deletions on non-coding transcription. As in the case of coding genes, overall transcription of non-coding transcripts is compromised upon DF deletion. Moreover, changes in Pol II occupancy in non-coding regions largely mirrored those in coding regions, particularly for NUT genes (compare Figs. 2 and S15), which suggests that initiation and early elongation is affected by the studied DFs similarly in both coding and non-coding regions. However, changes in non-coding 3’-proximal pausing of Pol II do not match what was observed for coding genes. First, we note that 3’ pausing in ncRNA regions is less apparent than in their coding counterparts, perhaps implying that they are regulated differently. Indeed, production of the 3’ ends of NUTs is controlled by a unique mechanism (Schulz et al. 2013). Perhaps a key distinction between transcription of coding and non-coding genes lies in the mechanisms controlling their respective terminations. Analyzing the effects of DFs at PAS of non-coding transcripts revealed that unlike in coding transcripts, deletions of the studied DFs did not lead to increased 3’ Pol II accumulation, implying that DFs interact differently with the PA machinery of coding and non-coding regions. Additionally, the lack of correlations between sense and antisense transcription for multiple ncRNA types and the insignificant effect of DF deletion on these correlations (results not shown) belies the possibility that regulation of antisense ncRNAs is the preferred mechanism by which DFs modulate coding transcription. Moreover, we found that genes which are highly sensitive to *XRN*1 deletion (i.e., the Xrn1 synthegradon) are less likely to have convergent anti-sense transcripts of any type (results not shown), suggesting that convergent ncRNAs are not the direct mechanism which mediates the effect of Xrn1 on coding regions.

Our data highlight additional distinctions between TFIID- and SAGA-dominated genes. On one hand, both the SAGA and TFIID complexes regulate transcription of nearly all genes (Bonnet et al. 2014; Baptista et al. 2017; Warfield et al. 2017), and their respective transcripts have comparable half-lives (Fig. 6D). Cumulative results, shown here and published by others (Tirosh et al. 2007; Kubik et al. 2015; de Jonge et al. 2017), suggest that the two classes are each characterized by distinct chromatin structure and different transcriptional plasticity. On the other hand, we found that deletion of *XRN*1, and to a lesser extent our other studied DFs, affects transcription of TFIID-dependent genes more than SAGA-dominated genes (Figs. 6A, 6B). Moreover, the binding of Xrn1, Lsm1, and Dcp2 to promoters occurs at different positions between the two classes, with SAGA-dominated genes having binding sites located further upstream of TSS than TFIID-dominated genes (Fig. 6E). In addition, we found that both Xrn1 and Ski2 bind SAGA-dominated gene transcripts more than TFIID-dominated gene transcripts; further, Xrn1 and Ski2 exhibit different preferences to the two classes of mRNAs (Fig. 6C). We thus suggest that a major difference between SAGA- and TFIID-dominated genes is not related to transcription or mRNA decay *per se*, but to the steady-state levels of their mRNA products. We propose that the two classes of genes differ in the buffering mechanisms controlling their mRNA levels, involving a linkage between RNA and chromatin binding features of some of the factors that we examine in this paper. In particular, we suggest that the deletions of *T AF*1 and *SPT* 3 compromise the cross talk between mRNA synthesis and decay for TFIID- and SAGA-dominated genes, respectively, leading to decreases in steady-state levels of specific mRNAs.

Previously, we classified genes based on their sensitivity to *XRN*1 deletion (Medina et al. 2014). Genes whose synthesis and transcript stability were highly sensitive to this deletion were labeled the “Xrn1-sythegradon”. Here we found that our *xrn*1Δ NET-seq data are consistent with our prior classifications, as Pol II occupancy fold changes nicely correlated with previously assigned responsiveness values (Fig. 1C, “*xrn*1Δ” panel). Our results further demonstrate that these scores agree well with NET-seq fold changes of *lsm*1Δ, *dhh*1Δ, and somewhat with *ccr*4Δ, but not with *rpb*4Δ (Fig. 1C). This raises the possibility that Dhh1 may also function together with Xrn1 and Lsm1 to link mRNA synthesis and decay, perhaps as part of the previously proposed complex containing the latter two proteins. In general, our analyses differentiate between two types of DFs. Type I comprises Xrn1, Lsm1, and Dhh1, the deletion of any one of which reduces 5’ Pol II occupancy and elongation rates of DF-stimulated genes but enhances Pol II pausing at PAS; Type II includes Ccr4 and Rpb4, whose deletions inhibit the release of Pol II from positions roughly 100 bp post-TSS or enhance elongation rates downstream of these locations (Figs. S6C, S6D). This distinction is further underlined by the aforementioned difference in the correlations between knockout-associated changes in NET-seq signals and Xrn1 responsiveness (Fig. 1C). Nevertheless, in contrast with the lack of correlation between DFs and the genes of the Rpd3S H4 deacetylation complex, we observed positive correlations among all studied DFs (Fig. 1B). We hence propose that they act similarly at the global level, akin to our previous suggestions for Xrn1, Lsm1, and Dcp2 (Haimovich et al. 2013).

In aggregate, our results point towards roles for Xrn1 and other DFs in facilitating the efficient elongation of Pol II early and, more clearly, late during transcription, potentially via control of pausing and backtracking. The identified functions of DFs further highlight the key roles of these processes in the regulation of transcription. In recent years, promoter-proximal pausing has been a focal point in the study of transcription of many metazoans; whether Pol II pausing plays a comparable key role at PAS and in transcription termination of *S. cerevisiae* and other organisms remains to be examined.

## Methods

### Data collection and pre-processing

We used the NET-seq protocol to measure the number of RNA polymerase II (Pol II) bound to DNA which are engaged in mRNA synthesis; experiments were performed as detailed in (Churchman and Weissman 2012). We consider the bound Pol II levels of 4973 genes in six genotypes comprising one control (in duplicate) and five mutant knockouts (*xrn*1Δ in duplicate). The data were pre-processed using cutadapt and prinseq (Martin 2011; Schmieder and Edwards 2011), and mapping was done via TopHat (Trapnell et al. 2009) with unique reads retained.

### Identification and interpretation of differentially transcribed genes

Normalization was performed by selecting genes whose productive transcription as measured by CDTA (*ccr*4Δ, *dhh*1Δ, and *lsm*1Δ) and GRO (*rpb*4Δ and *xrn*1Δ) was least perturbed (log2 FC between −0.25 and 0.25) by DF deletion (Sun et al. 2012; García-Martínez et al. 2004) as these are the genes for which we expect the least disruption in transcriptional processes. Each strain was then separately normalized to wild type samples by using the normalization procedure of DESeq2 restricted to the appropriate sets of genes (Love et al. 2014).

Standardized fold changes of Pol II levels in genes and ncRNAs were computed using DESeq2 (Love et al. 2014); these values account for the heteroskedastic mean-variance trend for gene Pol II counts. Gene ontology (GO) enrichment analyses among genes were performed using GOrilla in single-ranked list mode (Eden et al. 2007; Eden et al. 2009). Transcriptional efficiencies were computed by taking the log2 ratio of GRO to NET-seq reads (for whole genes) or BioGRO to NET-seq reads (for profiles).

### Metagene construction

Metagene densities (Figs. 2, S15) for entire transcript bodies were generated by re-scaling each transcript to a length of 1000 nt. We then aggregated reads across genes and applied lowess smoothing to create metagene profiles (Fig. 2). The metagene profiles of TSS-adjacent and PAS-adjacent sites instead display the average number of normalized reads in defined regions. Specifically, reads no more than 100 nt (500 nt) upstream (downstream) of an annotated TSS were incorporated for 5’ metagenes, while all reads within 150 nt of PAS were aggregated to give characteristic profiles for Pol II distributions near polyadenylation sites. We computed elongation efficiency metagenes in the WT and *xrn*1Δ strains by taking BioGRO reads in the same regions described above and smoothing the profiles for each gene. We then took the log2 of ratio of the smoothed BioGRO and NET-seq profiles for each gene and averaged these profiles across all genes.

### Application of mathematical model

Our mathematical analysis of initiation and elongation rates is based on the recently obtained analytical solutions (Erdmann-Pham et al. 2018) to the Totally Asymmetric Simple Exclusion Process (TASEP). Denoting the NET-seq Pol II metagene for a given sample at position *x* as *ρ*(*x*), site-specific elongation rates *λ*(*x*) were approximated as *λ*(*x*) *≈*1*/*(*ρ*(*x*)(1 *− ρ*(*x*))) after appropriate re-scaling of *ρ*(*x*) to lie in the interval (0, 1). Ideally, the re-scaling would be given by the number of cells in each sample such that *ρ*(*x*) is the probability of finding a Pol II at position *x*. As these numbers were unknown, a range of re-scalings were tried and the results were robust. It is worth noting that the actual initiation and elongation rates are not identifiable without the exact re-scaling; however, the ratio of initiation and elongation rates is, permitting our analysis.

## Data access

NET-seq data for WT, *xrn*1Δ, *dhh*1Δ, *lsm*1Δ, *ccr*4Δ, and *rpb*4Δ cells were generated as part of this project and accompany this release. External data sets are available as follows: Remaining NET-seq - GSE25107 (Churchman and Weissman 2011); CRAC - GSE46742 (Tuck and Tollervey 2013); RNA-seq FCs - GSE107841 (He et al. 2018); GRO - GSE29519, GSE57467 (Haimovich et al. 2013; García-Martínez et al. 2015); CDTA - E-MTAB-1525 (Sun et al. 2013); BioGRO - GSE58859 (Jordán-Pla et al. 2014; Jordán-Pla et al. 2016) and manuscript in progress; ChIP-exo - GSE44312 (Haimovich et al. 2013); mRNA half-lives accompany Medina et al. 2014.

## Acknowledgments

We thank Sebastián Chavez, Jośe E. Pérez-Ortín, and Antonio Jordán-Pla for their comments on our results and use of BioGRO data. We also thank Dan D. Erdmann-Pham for discussions regarding application of the computational model used for simulations. The research is supported in part by an Israel Science Foundation grant 1472/15, NIH training grant T32-HG000047, NIH grant R01-GM094402, and a Packard Fellowship for Science and Engineering. NY and YSS are Chan Zuckerberg Biohub Investigators.

## Author contributions

MC conceived the study and performed the experiments with some assistance from LSC. JdI performed the data pre-processing. JF performed the statistical and computational analyses with suggestions from MC, YSS, and NY. JF and MC wrote the manuscript. YSS, NY, and LSC critically read the manuscript.

